# Genomic insights into the recent population history of Mapuche Native Americans

**DOI:** 10.1101/2021.11.25.470066

**Authors:** Lucas Vicuña, Anastasia Mikhailova, Tomás Norambuena, Anna Ilina, Olga Klimenkova, Vladimir Shchur, Susana Eyheramendy

**Affiliations:** Faculty of Engineering and Sciences, Universidad Adolfo Ibáñez, Peñalolén, Santiago, Chile; Instituto Milenio de Investigación Sobre los Fundamentos de los Datos (IMFD), Santiago, Chile; International Laboratory of Statistical and Computational Genomics, HSE University, Russian Federation

**Keywords:** Mapuche, Native Americans, genetic history, adaptation

## Abstract

The last few years have witnessed an explosive generation of genomic data from ancient and modern Native American populations. These data shed light on key demographic shifts that occurred in geographically diverse territories of South America, such as the Andean highlands, Southern Patagonia and the Amazon basin. We used genomic data to study the recent population history of the Mapuche, who are the major Native population from the Southern Cone (Chile and Argentina). We found evidence of specific shared genetic ancestry between the Mapuche and ancient populations from Southern Patagonia, Central Chile and the Argentine Pampas. Despite previous evidence of cultural influence of Inca and Tiwanaku polities over the Mapuche, we did not find evidence of specific shared ancestry between them, nor with Amazonian groups. We estimated the effective population size dynamics of the Mapuche ancestral population during the last millennia, identifying a population bottleneck around 1650 AD, coinciding with a period of Spaniards’ invasions into the territory inhabited by the Mapuche. Finally, we show that admixed Chileans underwent post-admixture adaptation in their Mapuche subancestry component in genes related with lipid metabolism, suggesting adaptation to scarce food availability.

## 1 Introduction

Human presence in the American Continent dates back to at least 21000 years before present (BP), as confirmed by recent findings in North America [1, 2, 3]. According to the current understanding derived from archaeogenetics, the first Native American ancestral population diverged from East Asians, crossed Beringia 25000 −23000 BP and shortly split into two lineages 17000−14600 BP, called *Ancestral-A* (or *Southern Native American*) and *Ancestral-B* (or *Northern Native American*). While *Ancestral-B* originated eastern North American populations, *Ancestral-A* originated the lineage of the 12900−12700 BP Anzick-1 individual associated with the Clovis Culture and rapidly radiated to Central and South America [4, 5, 6].

The peopling of South America began at least 14000 before present (BP), as revealed by the archaeological site of Monte Verde in Southern Chile [7]. Around 3000 years later, an ancient lineage that split from *Ancestral-A* appears in Central Chile. This lineage is represented by an individual from Los Rieles (*Chile_LosRieles_10900BP*) [6] [8], has high genetic affinity with the Anzick-1 Clovis individual, and was almost completely replaced between 10900 − 9000 BP by a second *Ancestral-A* branch population. This latter lineage explains the genetic diversity of most present-day Native populations from Central and South America, with two exceptions: i) Central Andeans, who share an *Ancestral-A*-derived lineage related to 4900 BP individuals from the California Channel Islands (*USA_SanNicolas_4900BP*) [6]; and ii) some populations from the Amazon basin and the northwestern Coast of South America, who have a minor Australasian component [9] [10]. Around 5100 BP and 700 BP, there is evidence of high genetic affinity between individuals from Los Rieles (*Chile_LosRieles_5100BP*) and Conchali (*Chile_Conchali_700BP*), respectively, with modern Huilliche and Pehuenche [8], two Mapuche subpopulations from the Araucanía Region of southern Chile (**Figure 1**).

**Figure 1.**
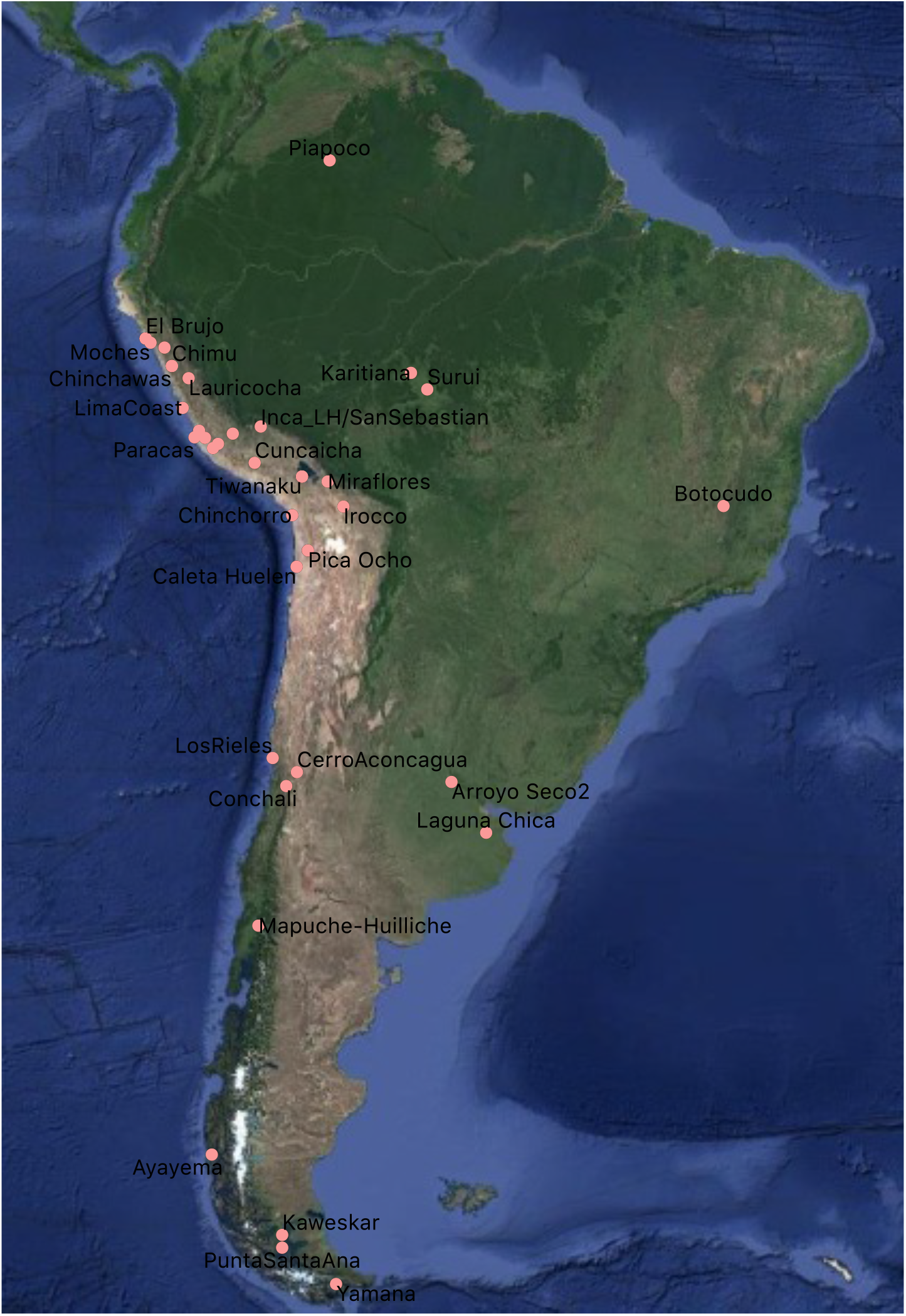
The geographic location of populations used in this study. Shown is the location of representative populations or archaeological sites from the Southern Cone, the Central Andes and the Amazon Basin. Some archaeological sites harbor 1 populations. (e.g., LosRieles harbors *Chile_LosRieles_5100BP* and *Chile_LosRieles_10900BP*). To avoid overfilling the map with text, we show only some populations (**Figure 2**). This map was done with GGI3.10 (https://www.qgis.org/en/site/)

The Mapuche (who self-denominated *Reche* at the time of contact with Europeans) are a Mapudungún-speaking Native group from Chile and Argentina, and the only extant Native population living in Central-Southern Chile. Based on their geographic location, the Mapuche have in general been divided in three main groups: Picunche (northern lowlands), Huilliche (southern lowlands) and Pehuenche (southeastern foothills of the Andes) [11]. The population history of the Mapuche has been debated. An hypothesis posited by historian Francisco Antonio Encina and taught in many Chilean schools today is that they arrived to the Araucanía Region from the Argentinian Pampas, crossing the Andes Mountains one or two centuries before the Inca invasions that took place between 1471 and 1532 AD [12] [13]; see also [14]). A more accepted hypothesis is that groups of hunter-gatherers lived in the Araucanía since ancient times and that one them imposed their culture over the others at least 1500 − 1400 BP [15]. Indeed, the Mapuche might be direct descendants of the pre-Columbian archaeological cultures Pitrén (100−1100 AD) and El Vergel (1100–1450 AD) of Araucanía [13]. The Picunche, who were part of the Aconcagua Culture (900−1450 AD), were conquered by the Inca Empire during their expansion to present-day North/Central Chile in 1438-1539 AD, as revealed by hundreds of Inca archaeological sites [13] and by historical records [15]. However, it is unknown whether these two populations underwent genetic admixture. Further, it is not clear whether southern Mapuche, namely, Huilliche and Pehuenche, were culturally or genetically influenced by the Inca [13].

It has also been suggested that the Mapuche had pre-Columbian cultural exchange with the Tiwanaku Culture (∼ 600 − 1100 AD) from the Central Andes as well as with Amazonian groups. Dillehay et al., 2007 [16] hypothesized that the Mapuche society underwent a series of cultural changes following the collapse of the Tiwanaku empire on ∼1100 − 1300 AD, causing a southward migration wave that would have reached the Araucanía Region. Similarly, the authors hypothesized that ancient populations from Araucanía may have been culturally influenced by groups from the southern Amazon basin. Both cultures have similar ceramic traditions with apparent roots in central Chilean or northwest Argentine cultures [16]. Despite these observations, it is not known whether the Mapuche share specific genetic ancestry with Andean or Amazonian groups.

During the 15^*th*^ and 16^*th*^ centuries, the Mapuche resisted the invasions of the Inca Empire [17] and the Spanish Kingdom [18]. The arrival of Spaniards brought along infectious diseases that caused severe epidemics among the Mapuche, such as typhoid fever (1558 AD) and smallpox (1563 AD) [15]. Arguably, wars and diseases resulted in many deaths among the Mapuche [15]. The arrival of Spaniards also resulted in their admixture with the Mapuche, originating modern Chileans centuries later. Possibly, Chileans inherited adaptation signals from their Native American ancestors, to cope with selective pressures such as infectious diseases, food scarcity and extreme climates, similar to other populations with Native American ancestry [19, 20, 21, 22, 23, 24, 25].

The present study had three aims: i) Looking for specific shared ancestry between the Mapuche and populations from the Southern Cone, the Central Andes and the Amazon Basin; ii) Estimating the effective population size (N_*e*_) trajectory of the Mapuche lineage during the last millennia; and iii) Detecting Mapuche-specific signatures of natural selection inherited by modern admixed Chileans.

## 2 Results

Here we present several results which put together the whole picture of the Mapuche specific history. We identified specific shared ancestry events between the Mapuche and ancient populations from the Southern Cone. We confirmed that the Mapuche trace most of their ancestry from a lineage present in Central Chile at least 700 BP [8] (Aconcagua Culture [13]), and probably before 5100 BP. We also identified events of gene flow with Southern Patagonian groups. Despite previous observations of cultural exchange between important Andean polities with the Mapuche (such as Inca and Tiwanaku) [16, 13], we did not find genetic exchange between them, nor with Amazonian groups. Our analysis demonstrates that there is an agreement between the Mapuche effective population size decline inferred from genetic data and a period of diseases and wars caused by foreign invasions into Mapuche territory between the 15^*th*^ and 17^*th*^ centuries [15, 17]. Finally, we were able to capture Mapuche-specific adaptation signals inherited by admixed Chileans. These results provide a comprehensive insight in the history of the Mapuche population, which is the major Native American population in Chile and Argentina.

### 2.1 Evidence of specific shared ancestry between the Mapuche and ancient populations from the Southern Cone

We estimated genetic affinity between the Mapuche and Native South American populations from distinct geographical regions. For this, we used a set of 124, 470 variants that fulfilled two conditions: i) They have < 0.01 non-Native American local ancestry mean in our sample of 11 Mapuche-Huilliche individuals; and ii) They intersect between the Mapuche-Huilliche dataset and the 1240K HO dataset of the Allen Ancient DNA Resource (AADR) [26] (see Methods for details).

We first analyzed whether ancient populations from the Southern Cone region of South America were ancestors of the Mapuche (Southern Cone was broadly defined as the regions from Chile and Argentina located at the same or lower latitudes than the archaeological site of Los Rieles, see **Figure 1**). We used 11 individuals belonging to the Huilliche branch of the Mapuche (Mapuche-Huilliche). We computed D(Mbuti, Ancient Southern Cone; Modern South America, Mapuche) statistic, iterating over different combinations of Ancient Southern Cone and Modern South American populations. Modern was defined as ≤ 100 BP. We found consistent evidence of allele sharing between *Chile_Conchali_700BP* (700−900 year-old individuals from Central Chile) and the Mapuche-Huilliche, as revealed by Z-scores > 3, in 7 independent significant tests (**Supplementary File 1A** and **Figure 2A**), confirming previous findings [8]. We also found evidence of allele sharing between Mapuche-Huilliche and *Chile_LosRieles_5100BP* (2 significant tests), *Argentina_ArroyoSeco2_7700BP* (1 significant test), *Chile_WesternArchipelago_Kaweskar_1200BP* (1 significant test), and *Chile Yamana_BeagleChannel_800BP* (2 significant tests), relative to modern Andean and/or Amazonian populations (**Figure 2A**).

**Figure 2.**
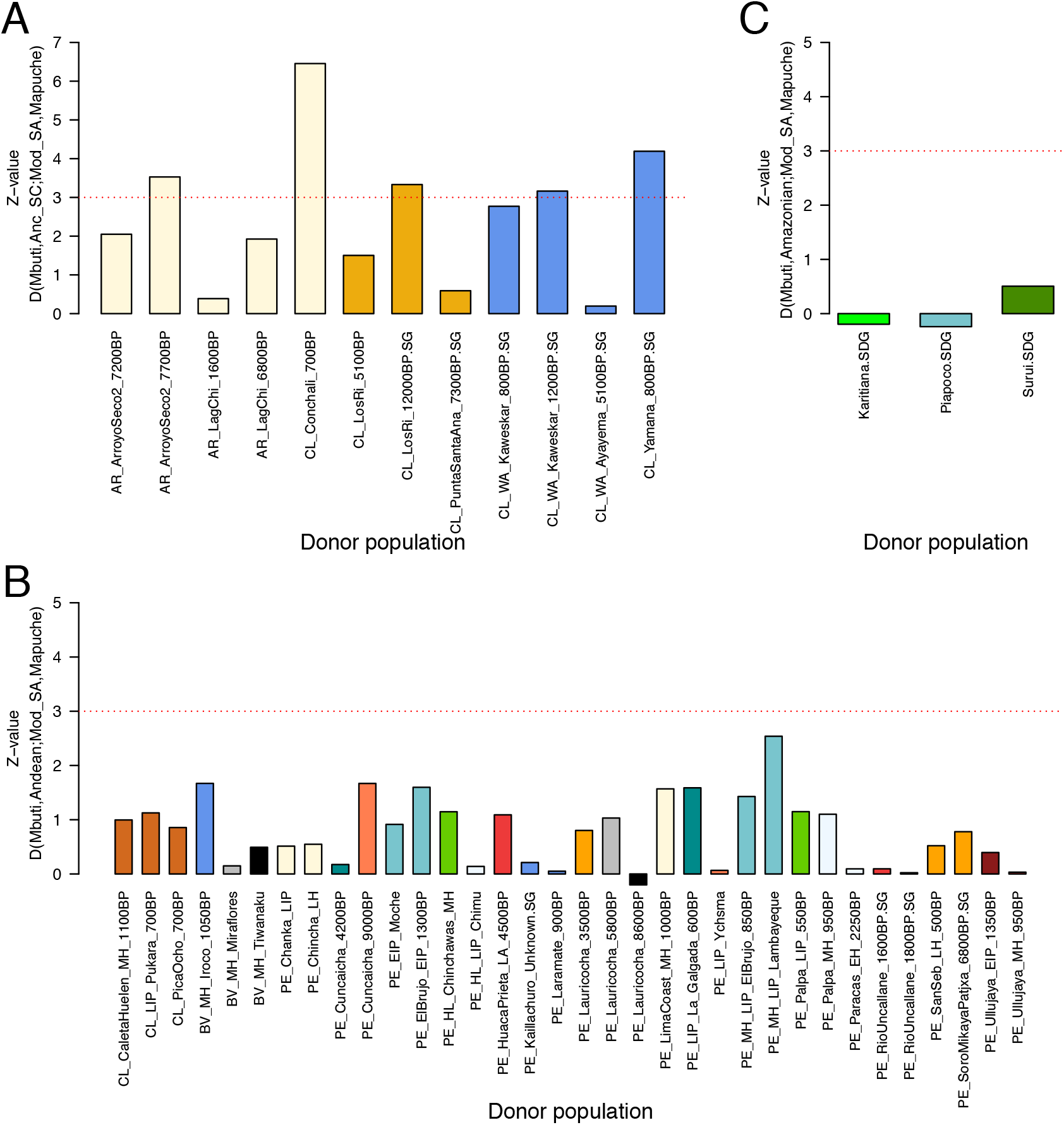
Genetic a_nity between Mapuche-Huilliche and Southern Native American populations. Distribution of Z-values (y-axis) for D statistic tests between a donor population and the Mapuche-Huilliche. **A**. The donor population is Ancient Southern Cone (Anc SC). **B**. The donor population is Central Andean. **C**. The donor population is Amazonian. The red dotted horizontal line represents the significance threshold of Z = 3. For results of the same population but different sequencing technologies (e.g., Karitiana.DG vs. Katitiana.SG [26]), the figure shows the worst Z-value. For each donor population/Mapuche-Huilliche pair, the figure shows the result with the highest Z-value. All remaining test results are found in `Supplementary File 1A-C`. With a few exceptions, the figure shows tests with Z> 0. Due to space limits, we show short versions of the original populations’ names. Conversions between the original (long) names and short names are found in `Supplementary File 1D`.

**Supplementary File 1A** shows the tested populations and the corresponding Z-scores. We evaluated whether close genetic relatedness between Mapuche-Huilliche individuals could affect the D-statistic results. Thus, we performed the same tests described before but keeping Mapuche-Huilliche individuals without relatedness of first and second degree, obtaining a set of 6 individuals. The results were almost identical (data not shown). **Figure 1** shows the geographic location of populations from representative regions used in this study.

We used `qpGraph` [27] to model the phylogenetic relations between the Mapuche-Hulliche and other ancient as well as modern Southern Cone populations. Based on our D-statistic results and on previous results [8] [6] [28], we started our phylogenetic tree including the Anzick1 Clovis individual (*USA_Anzra_SG*), *Argentina_ArroyoSeco2_7700BP, Chile_LosRieles 5100BP* and *Chile-Conchali_700BP*. We used *Mbuti* as outgroup and included *Aymara* as a representative modern descendant of the California Channel Islands-related Native American branch population. Later, we included ancient populations from Southern Patagonia. We obtained a model tree by fitting the Mapuche-Huilliche as a mixture of a lineage from Central Chile (related to *Chile Conchali_700BP* ; 97%) and an ancient lineage from Southern Patagonia (3%; related to *Chile WesternArchipelago_Kaweskar 1200BP*.*SG*) (**Figure 3**).

**Figure 3.**
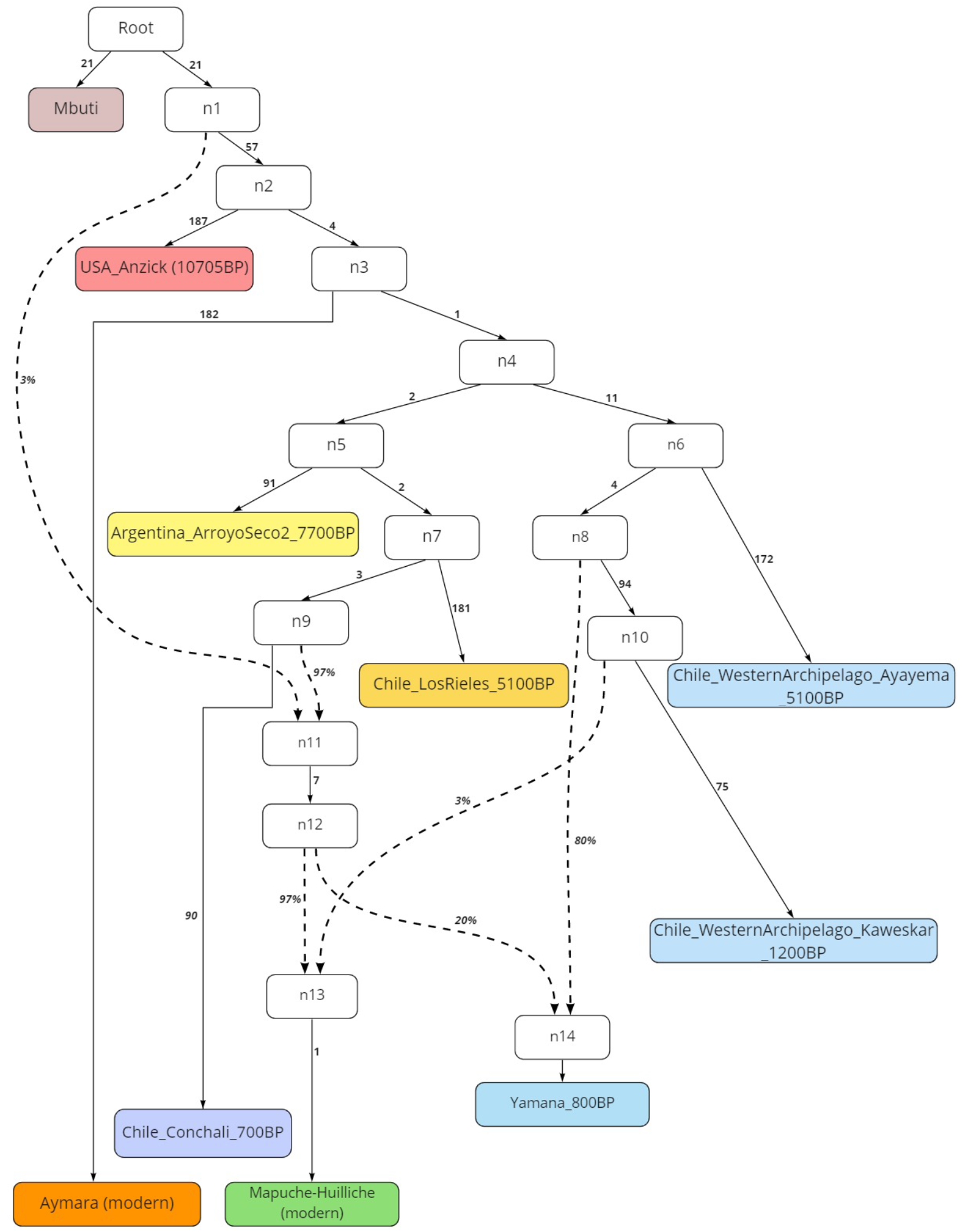
Admixture graph relating the Mapuche-Huilliche with ancient Southern Cone populations. Numbers along branches represent units of shared genetic drift. Nodes labeled with n1, n2, etc., represent unknown populations necessary to fit the model. Percentages indicate admixture proportions. Z-score o8f the worst *f* 4(Anzick1, Aymara, *Argentina_ArroyoSeco_700BP, Chile_Conchali_700BP*) is 2.585.. This graph was made with Miro (https://miro.com/)

### 2.2 No evidence of specific shared ancestry between the Mapuche with Central Andean or Amazonian populations

We tested for specific shared ancestry between the Mapuche-Huilliche and Central Andean populations, such as Inca and Tiwanaku. We included Inca individuals from different locations as well as Tiwanaku individuals from time periods covering the whole Tiwanaku civilization (see Discussion) [28]. We computed D(Mbuti, Ancient Central Andean; Modern South America, Mapuche) statistic, by iterating over all combinations of ancient Central Andean populations (**Figure 2B**) and modern South American populations. We did not find evidence of specific shared ancestry among these groups (**Figure 2B**). **Supplementary File 1B** shows the tested populations and the corresponding Z-scores.

We modeled the phylogenetic relations between the Mapuche-Huilliche and Central Andean populations, similarly as before. We included Tiwanaku (*Bolivia_MH_Tiwanaku*), Inca individuals (*Peru_SanSebastian_LH_500BP* and *Peru-LH_Inca*), as well as an older individual from a nearby archaeological site (*Peru_Laramate_900BP*). We obtained a model fitting the Mapuche-Huilliche as a mixture of *Chile_Conchali_700BP* (66%) and an undefined Native American branch (34%) (**Figure 4**).

**Figure 4.**
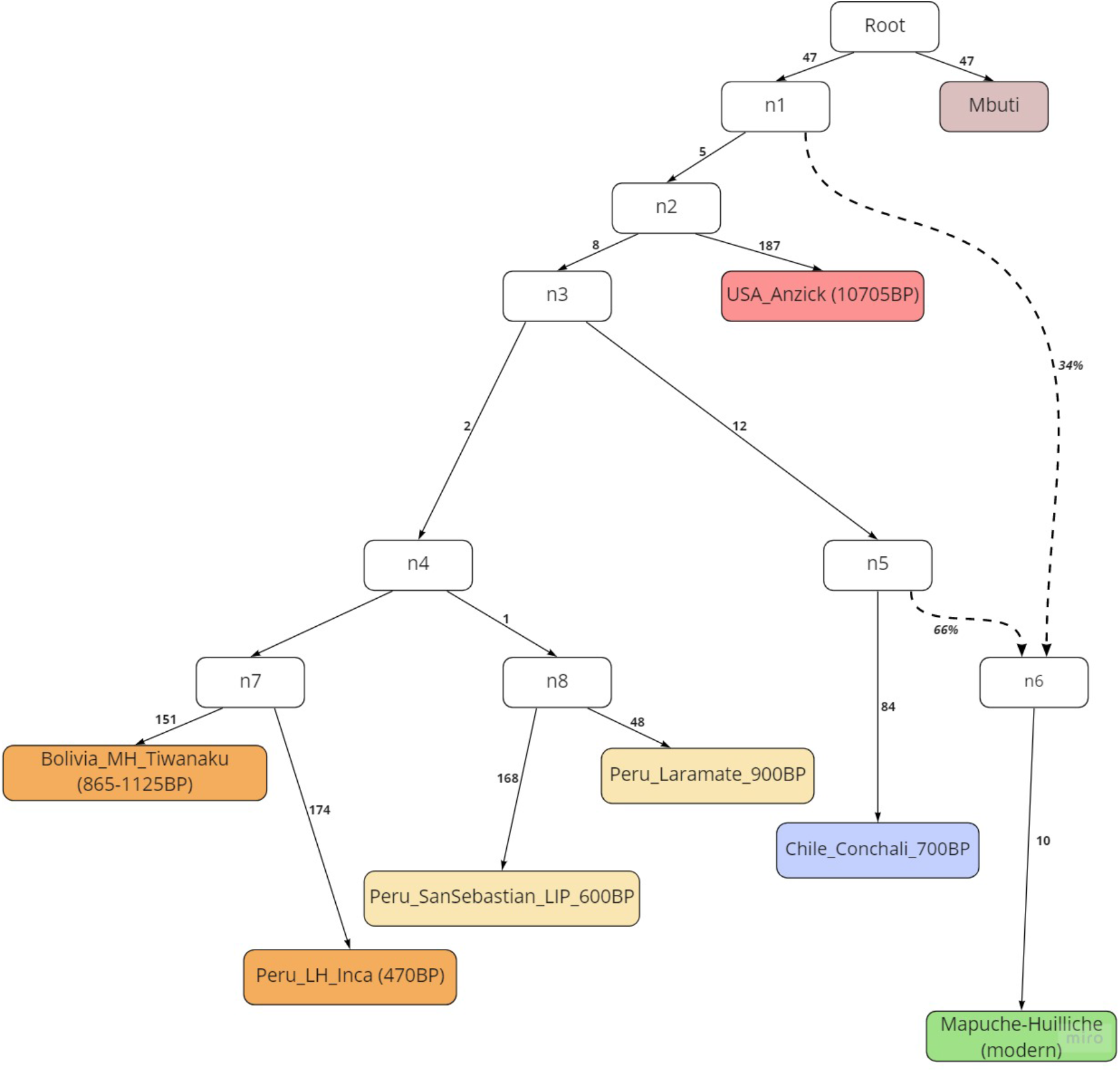
Phylogenetic tree relating the Mapuche-Huilliche with ancient Central Andean populations. Numbers along branches represent units of shared genetic drift. Nodes labeled with n1, n2, etc., represent unknown populations necessary to fit the model. Percentages indicate admixture proportions. *Z*-score of the worst *f* 4(*Peru_LH_Inca, Peru_Laramate_900BP, Chile_Conchali_700BP, Peru_SanSebastian LIP 600BP*) is 2.166.. This graph was made with Miro (https://miro.com/).

Our D-statistic results also show that there is specific shared ancestry between modern Amazonian and ancient Central Andean populations relative to the Mapuche, as revealed by Z-scores < -3 (e.g., *Karitiana* vs. *Peru_Chincha_LH, Peru_Ullujaya_MH_950BP, Peru_LimaCoast_MH 1000BP* and *Peru_MH LIP_La-mbayeque*) (**Supplementary File 1B**)

We tested for genetic affinity between the Mapuche-Huilliche and the following modern populations from the Amazon Basin (hereafter referred as “Amazonian”) for which there is public genomic data: *Surui, Piapoco* and *Karitiana*. We computed D(Mbuti, Amazonian; Modern South America; Mapuche) statistic, iterating over modern Amazonian and modern South American populations. We did not find evidence of significant allele sharing between Amazonian groups and the Mapuche-Huilliche (**Figure 2C** and **Supplementary File 1C**).

### 2.3 Mapuche effective population size

We used single nucleotide polymorphism (SNP) data from modern admixed Chileans [29] to estimate how the effective population size N_e_ of Mapuche’s lineage changed over the last millennia. We implemented a methodological approach based on the relative length of Identity-by-descent (IBD) segments of specific ancestries, which is suitable for SNP array data [30]. Specifically, the Chilean individuals used in our study have on average 0.438 Mapuche Native American, 0.026 Aymara Native American, 0.521 European and 0.015 African mean global ancestry proportions [31]. In order to analyze N_e_ changes that are specific to the Mapuche, we excluded 592 admixed individuals with 0.01 global Aymara ancestry, obtaining a final dataset of 312 individuals. Based on segments of at least 2 centiMorgan genetic length, this method is able to accurately estimate N_e_ within the last hundreds of generations. Briefly, we used the refinedIBD program [32] to detect such segments, RFMix [33] to estimate the (local) Native American ancestry of each SNP (haplotype) and IBDNe [34] to estimate ancestry-specific N_e_. We observed that the Mapuche ancestral component underwent a population bottleneck that reached a minimum of N_e_ = 10, 300 at 13 generations ago (year ∼1650 AD) compared to N_e_ = 27, 200 at 29 generations ago (year ∼1208 AD). Since the bottleneck, the N_e_ showed a steady increase until the present time (**Figure 5**).

**Figure 5.**
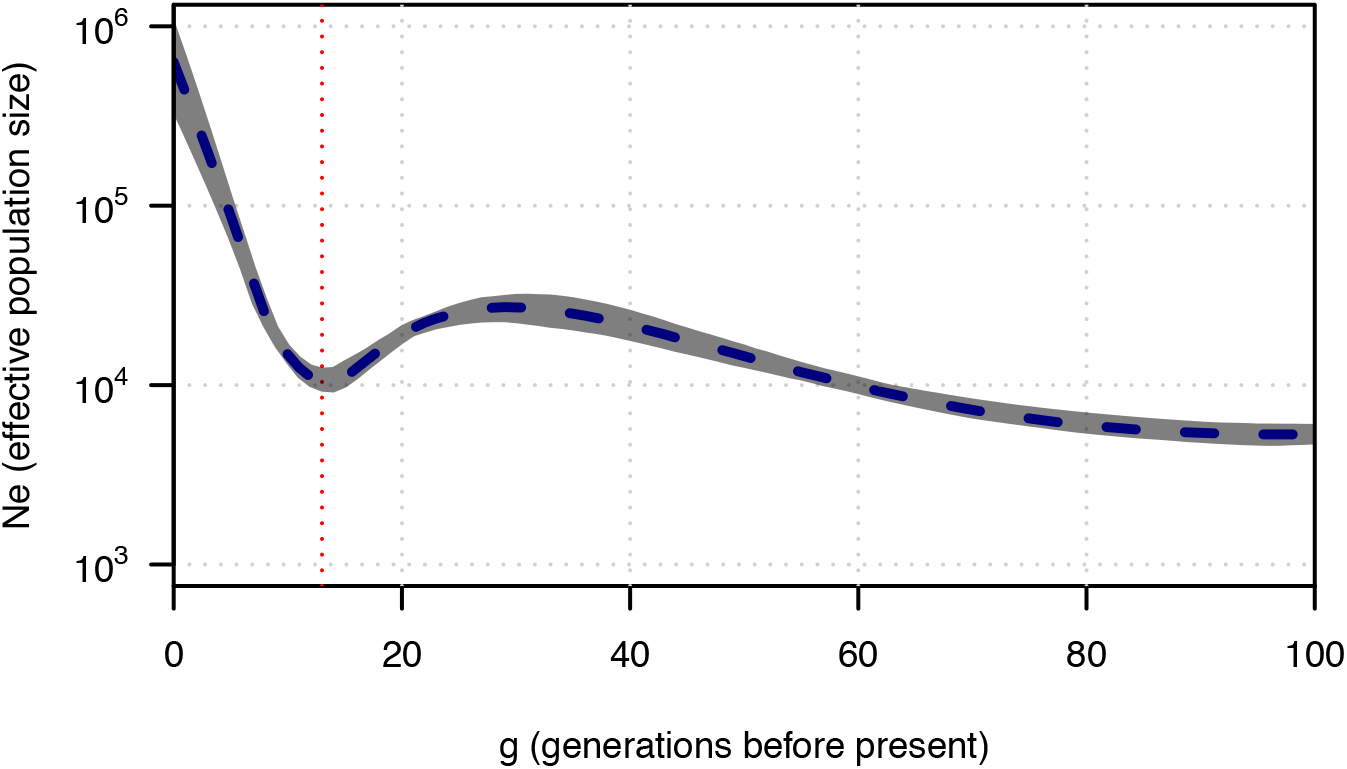
Temporal trajectory of Mapuche-specific N_e_. N_e_ trajectory (shown on log-scale) as a function of generations (g) before present. The dotted vertical line represents the recent (≤ 50 g) minimum N_e_ in the Mapuche ancestral component, corresponding to 13 generations ago. The grey area represents the 95% bootstrap confidence intervals. The plot is based on the analysis of 312 admixed individuals with Mapuche, European and African ancestries.

### 2.4 Post-admixture selection in the Mapuche component of admixed Chileans

In order to identify genomic regions enriched in Mapuche ancestry but not Aymara ancestry, we implemented the following strategy. 1) We performed *T* -tests to detect extreme deviations in the Native American local ancestry proportion over the genome-wide Native American mean in the full set of 904 admixed individuals. We identified 2, 247 variants crossing the genome-wide threshold of *P* < 10^−5^ recommended for recently-admixed populations [35]. 2) To minimize potential confounding effects produced by Aymara ancestry, we only considered the 312 individuals with ≤0.01 Aymara Native American ancestry. 3) We performed the same *T* -tests on the subset of 312 individuals, identifying 467 variants with *P* < 10^−5^. 4) To avoid a possible ascertainment bias introduced by selecting only individuals with Mapuche ancestry, we intersected the SNPs with significant deviations in the set of 904 individuals and in the set of 312 individuals. This resulted in a total of 65 SNPs. All of these SNPs achieved an association *P* -value of *P* = 2.4 × 10^−6^ and they are located within five genes mapping close together on chromosome 16, namely, *CPNE2, FAM192A, RSPRY1, PLLP* and *ARL2BP*. **Figure 6** shows genome-wide deviations in Native American ancestry in the subset of 312 individuals enriched with Mapuche ancestry, highlighting genes associated with the 65 significant variants that intersected between the two sets of individuals. **Supplementary Table 1** shows the 65 SNPs and their variant annotations.

**Figure 6.**
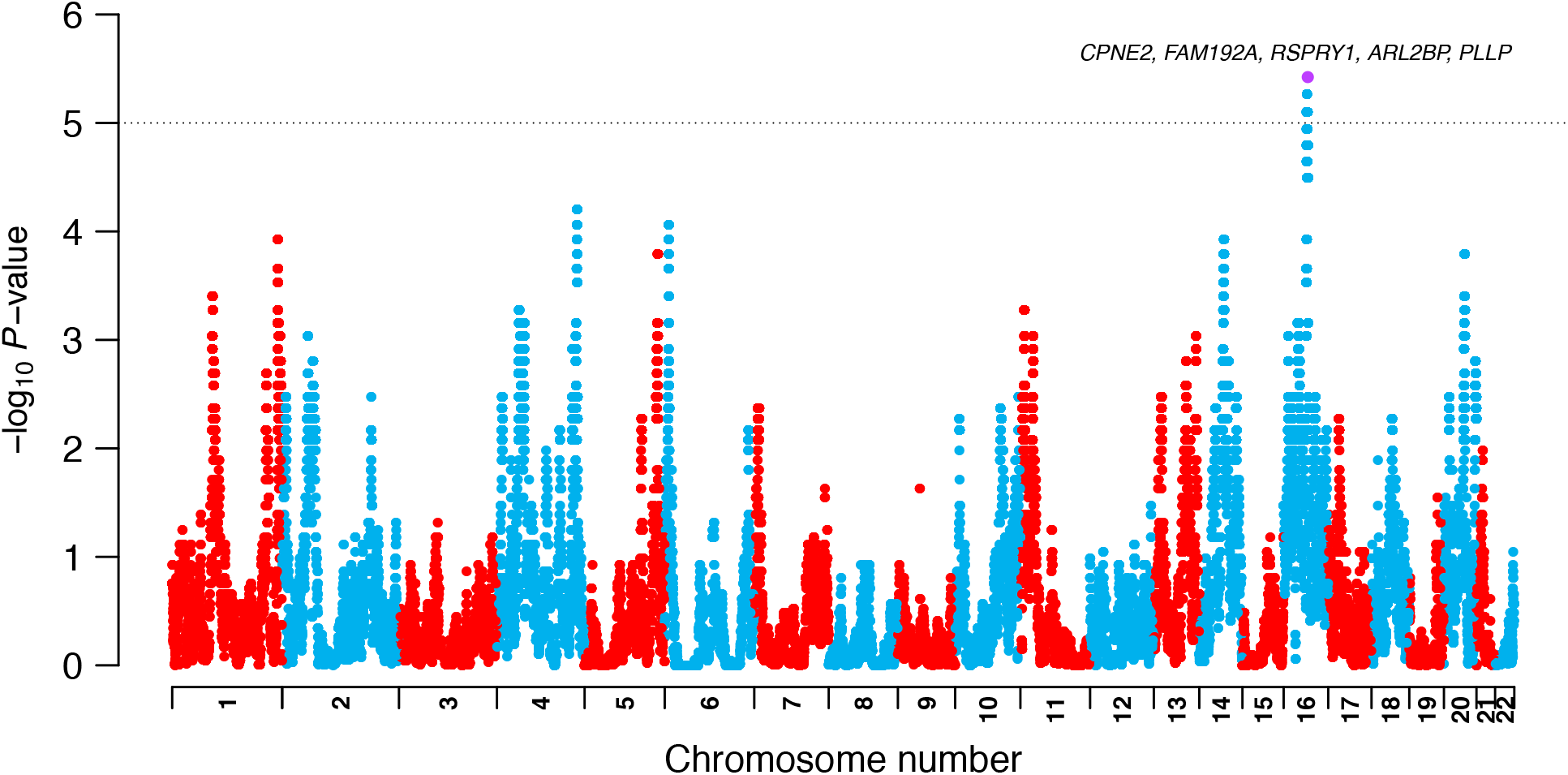
Genome wide deviations in Mapuche ancestry among Chileans. Deviations in Native American local ancestry over the genome-wide Native American local ancestry mean, expressed as the −log_10_(*T* -test *P* -value) of deviations. The dotted horizontal line represents the cutoff threshold of *P* < 10^−5^ for deviations considered significant [35]. Highlighted in violet are genes associated both with selected variants in the set of 312 individuals as well as in the 904 individuals.

In order to rule out that these significant deviations were caused by genetic drift, we performed neutral simulations of local ancestry (see Methods for details). We set a single admixture event between Native Americans, Europeans and Africans in proportions 0.46, 0.52 and 0.015, respectively, mimicking the real data. Admixture occurred 10 generations ago, which was the time where the main pulse of admixture occurred in admixed Chileans [36]. The effective population size of admixed Chileans was set to (N_e_) = 6, 000, based on previous estimates [37]. We simulated sample sizes of *N* = 156, 312, 624 and 904. For each *N*, we calculated *T* -test *P* -values for the SNP with a maximal deviation in Mapuche ancestry, obtaining *P* = 0.00877, 0.00103, 0.00214, 0.00006 for *N* = 156, 312, 624, 904 respectively. Thus, no simulated SNP achieved the significance threshold of *P* < 10^−5^ recommended for this test in recently admixed populations [35], ruling out that the significant deviations observed in the real SNPs were caused by drift.

## 3 Discussion

In this study, we tested hypotheses of specific shared ancestry between the Mapuche-Huilliche from Southern Chile and geographically distant populations from Native South America (**Figure 1**). We found consistent evidence of significant allele sharing between the Mapuche-Huilliche and lineages represented by *Chile_Conchali_700BP* and *Chile_LosRieles_5100BP* in Central Chile, confirming previous findings [8]. *Chile_Conchali_700BP* contributed most of Mapuche-Huilliche’s ancestry. This is in agreement with anthropological evidence showing a cultural influence from groups of Central Chile over groups of Araucanía. For instance, there are clear similarities in the pottery style (e.g., asimetric crocks called *jarros pato* in Spanish) between the Llolleo Culture (1800 − 1200 BP) of Central Chile and the Mapuche as well as the Pitrén Culture of Southern Chile (100 − 1100 AD) [13]. We also found significant allele sharing between the Mapuche-Huilliche and a lineage represented by individuals of *Argentina_ArroyoSeco2_7700BP* found in a site close to the northwestern Atlantic Coast of Argentina (**Figure 1**). Further, we found significant allele sharing between the Mapuche-Huilliche and populations from Southern Patagonia, namely, *Chile_WesternArchipelago_Kaweskar_1200BP* and *Chile Yamana_BeagleChannel_800BP*. Together, these observations support the hypothesis that the Mapuche-Huilliche are mainly the descendants of a lineage present in Central Chile at least 700 BP, and probably before 5100 BP, with some gene flow from Southern Patagonian lineages.

It is well known that the Inca Empire (or *Tawantisuyo* in Quechua language) (1438-1539 AD) occupied a big fraction of present-day Central Chile. This region was included in the so-called *Collasuyo* Region of the Empire and was populated by the Mapuche-Picunche. This region harbors hundreds of Inca archaeological sites [13], including *pukara* (fortresses), cemeteries, agricultural and hydraulic complexes, roads, among others. Many of such sites contain Inca cultural objects of personal use, such as pottery, clothes, *quipu* (textile system for counting), knives, *pichcas* (miniature games) and others [13], indicating a clear influence over local cultures. Different kinds of evidence indicate an influence of the Inca over the Picunche. The latter coexisted with the Inca at the time of Europeans’ arrival [15]. Also, the center of Chile’s capital city Santiago, which lies within former Picunche territory, was once an Inca settlement [38]. The presence and cultural influence of the Inca in the Araucanía Region, where the Mapuche-Huilliche lived, is more contentious, in part due to the preservation of a few distinct cultural objects. For instance, the pottery style of local cultures contemporary to the Inca, such as El Vergel, does not indicate a clear Inca influence [13] (but see [39]). We did not find evidence of significant allele sharing between the Mapuche-Huilliche and the Inca, represented by a *Peru LH Inca* individual from 470 BP found close to Ayacucho, Perú, and by 500 BP individuals from San Sebastián, close to Cuzco, Perú [28]. One explanation would be that the Mapuche-Huilliche did not admix with the Inca. Another possibility is that the ancestors of the 11 Mapuche-Huilliche individuals used in this study lived in relative isolation until present, since they inhabit the Huapi Island, located in the Ranco Lake [40]. We also did not find significant genetic affinity between the Mapuche-Huilliche and the lineage of an Inca boy found in the Aconcagua Mountain [5] (**Supplementary File 1A**), which is located at the eastern border of former Picunche’s territory. It is likely that this Inca boy was brought from the Central Coast of Peru [41] [5] and thus does not represent an Inca-like lineage. Of note, the Aconcagua Mountain is located ∼100km away from the Conchalí archaeological site in Central Chile (*Chile_Conchali_700BP*) (**Figure 1**), which was also Picunche’s territory, but the *Chile_Conchali_700BP* lineage does have high genetic affinity with the Inca boy [28].

A previous study hypothesized that the Mapuche had cultural exchange with the Tiwanaku [16], a powerful preincaic empire that flourished from around 600 to 1100 AD around the Titicaca Lake in present-day Bolivia and Perú [42]. For reasons not completely understood, the Tiwanaku Empire collapsed by ∼1100 − 1300 AD, possibly leading to groups migrating southwards towards Central/South Chile [16]. Raised fields dated ∼1000 AD were found in the Araucanía Region where the Mapuche-Huilliche lived [16]. These fields have similar technological and typological features as those found 3500 − 2500 km to the north along the Peruvian Coast, at the lowlands of Northeast Bolivia and at the Lake Titicaca Basin of Northwest Bolivia [16]. The sudden appearance of such fields after AD 1000 suggests that they might have been introduced by the Tiwanaku [16]. Despite these observations, we did not find evidence of specific shared ancestry between the Mapuche-Huilliche and individuals from different time periods of the Tiwanaku Empire (1125BP, 990BP, 900BP and 865BP [28] [26]). Neither did we find evidence of shared ancestry with other Central Andean populations. However, it is possible the there was a direct or indirect cultural influence of some tested Central Andean populations over the Mapuche-Huilliche, but without gene flow.

Dillehay *et al*., 2007 [16] also hypothesized that a cultural exchange occurred between Mapuche and Amazonian groups. This is based on similar ceramic traditions, funeral practices [43, 44], shared genetic markers and linguistic similarities between Mapudungún and languages spoken by southern Amazonian groups such as Arawak, Tacanan and Guarani [16] [13]. However, we did not find evidence of specific shared ancestry between Amazonian populations and the Mapuche-Huilliche. Nevertheless, we cannot completely rule out shared ancestry, since we included a subset of three Amazonian populations. Future research will be needed to test for genetic affinity with other Amazonian groups.

Our results show that the Mapuche’s N_*e*_ exhibited a steady decline starting 30 generations ago, and reaching a minimum at 13 generations ago (year 1650 AD). This timing is relatively similar to a previous study’s estimate that modeled an abrupt population bottleneck for the Mapuche-Huilliche at ∼ 325 BP (year ∼ 1700 AD) using imputed SNP array data [22]. We estimated a gradual and milder decrease in Mapuche’s N_*e*_ from N_*e*_ ∼ 27, 200 (29 generation ago) to N_*e*_ *∼*10, 300 (13 generations ago). This decrease was 62.1%, thus milder than the 94% reduction estimated in the aforementioned study. The decline in Mapuche’s N_*e*_ is similar in magnitude as that underwent by the Native American component of Ecuatorians and Nicaraguans, lower to that exhibited by Mexicans [30] and higher than Native Americans from Brazil [45]. One possible cause of the decline in the Mapuche’s population size is war. Between 1485 and 1493 AD, the Mapuche-Picunche fought the Battle of the Maule against the Inca in response to the southward expansion of this empire into present Central Chile (unknown date between 1471 and 1532 AD) [17, 46]. It is likely that this battle resulted in many casualties among the Mapuche-Picunche, while the Huilliche and the Pehuenche would have been mostly unaffected due to their distant location. Later on, the Mapuche resisted the Spaniards’ invasions between 1546 -1662 AD during the long Arauco War fought in the Araucanía Region, which ended with the victory of the Mapuche [18]. The number of deaths among the Mapuche is unknown. A second cause of Mapuche’s population decline were epidemics spread by Europeans. According to historical estimates, epidemics of typhoid fever (1558 AD) and smallpox (1563 AD) would have killed 30% and 10% of the total population of 1, 000, 000 individuals, respectively. The most affected group would have been the Picunche, who had more contact with Spaniards [15].

Our ancestry enrichment test detected significant deviations in the local Mapuche ancestry proportion in the genes *CPNE2, FAM192A, RSPRY1, PLLP* and *ARL2BP*. Interestingly, most of these genes have been associated with lipid-related traits in genome-wide association (GWA) studies. *CPNE2* variants are associated with high density lipoprotein (HDL) cholesterol levels (*P* = 5 × 10^−12^) [47]. *RSPRY1* variants are associated with HDL cholesterol levels (*P* = 4 × 10^−42^) and Apolipoprotein A1 levels (*P* = 3 × 10^−30^) [47]. *PLLP* variants are associated with HDL cholesterol levels (*P* = 6 × 10^−12^) [48]. In addition, *PLLP* variants associated with traits related to red blood cells, such as mean corpuscular hemoglobin concentration (*P* = 2 × 10^−12^), hematocrit (*P* = 2 × 10^−14^), hemoglobin concentration (*P* = 1 × 10^−11^) and red blood cell count (*P* = 1 × 10^−12^) [49]. *FAM192A* and *ARL2BP* have not been GWAS-associated to any trait. Extreme deviations of local Mapuche ancestry in genes involved in lipid metabolism might be due to post-admixture selection in response to scarse food availability among the first admixed Chileans. This may have resulted in modern Chileans tending to accumulate more fat in their body in a time where food scarcity is not a problem (see “thrifty hypothesis” [50]). Indeed, present-day Mapuche and admixed Chileans have a high incidence of lipid-related disorders such as insulin resistance, obesity, cholesterol gallstones and metabolic syndrome [51, 52].

In summary, our results provide new insights into the recent population history of the Mapuche as well as adaptations underwent by them, adding more pieces to the puzzle of Native American’s genetic history.

## 4 Subjects and Methods

### 4.1 Samples

We used genomic data from 11 Mapuche-Huilliche individuals [40]. We used SNP array data from 904 admixed Chilean individuals [31] from the “Growth and Obesity Chilean Cohort Study” (GOCS) [29]. We obtained curated genomic data of ancient and modern Native American populations from the Allen Ancient DNA Resource (1240K HO dataset V44.3), typed at 597, 573 sites [26]. **Supplementary Table 1D** lists the populations used in this study, which have been previously published [53, 54, 4, 55, 56, 6, 5, 22, 57, 58, 59].

### 4.2 Local Ancestry Estimation

We used RFMix [33] to infer the local ancestry of haplotypes in the Mapuche-Huilliche and in the admixed Chilean individuals. We used populations from the 1000 Genomes Project [60] as references for Native Americans, Europeans and Africans. As source for Native American ancestry we used Peruvians (PEL) with >95% Native American ancestry (n=29). For African ancestry we used Yoruba in Ibadan, Nigeria (YRI, n=108). For European ancestry we used Iberian populations in Spain (IBS, n = 107), which best resemble the Spaniard ancestral component of populations from modern Chile [61]. RFMix requires phased haplotypes, which were inferred with Beagle v.5 [62], using the HapMap37 human genome build 37 recombination map. We ran RFMix using the following parameters recommended in the manual: `PopPhased -n 2 -w 0.2 --fb 1`. We inferred local ancestry at 4, 596, 314 variants among the Mapuche-Huilliche.

### 4.3 Global Ancestry Estimation

Global ancestry proportions of Mapuche-Huilliche and admixed Chileans were estimated with Admixture 1.3 [63], similarly as described previously [31]. For the African and European ancestry, we used the same reference populations mentioned in the previous section. For the Native American ancestry, we used the 11 Mapuche-Huilliche individuals [40], 64 Aymara individuals [64, 22] and the 29 Peruvian (PEL) individuals with > 95% Native American ancestry.

### 4.4 Deviations of local ancestry

The mean local ancestry at each SNP was estimated using a published R script [37]. The Native American proportion at each SNP was compared with the Native American genome-wide mean using one-tailed *T* -tests, as described previously [37]. Variants achieving a significance threshold of *P* <10^−5^ were considered to be under post-admixture selection [35].

### 4.5 Identification of unadmixed Native American haplotypes

The Mapuche-Huilliche individuals from our dataset have 4 − 8% non-Native American global ancestry proportions, mainly of European origin [40]. In order to obtain unadmixed Native genomic segments, we excluded variants showing > 0.01 mean European or African local ancestries, obtaining 1, 958, 132 variants. Among them, we obtained a total of 124, 470 variants that intersected between the Mapuche and 1240K HO datasets. We used KING software [65] to detect and exclude five individuals with second degree relatedness, namely, GS000011194-ASM, GS000011195-ASM, GS000011196-ASM, GS000011201-ASM and GS000012210-ASM).

### 4.6 D-statistics

Before computing D-statistics with ADMIXTOOLS [27], we solved some problems of format conversions related to the `convertf` program. When converting from EIGENSTRAT to Plink binary, the resulting `fam` file had ID numbers in column 1 and population labels in column 6 (e.g. 13199 Mapuche:GS000011196-ASM 0 0 2 Mapuche). Because the Plink `fam` format requires the Family ID (i.e. population ID) to be in the first column, columns 1 and 6 in the `fam` file were switched. Using Plink 1.9, 124, 470 SNPs intersecting between the Mapuche and 1240K HO datasets were extracted. The 1240K HO dataset had individuals with long IDs that killed the `convertf` process when converting from Plink binary to EIGENSTRAT format. Thus, long individual names were coded with shorter names (**Supplementary File 1D**). Using `--keep`, we kept individuals and populations used in our further analyses. Plink binary files were back-converted to EIGENSTRAT using `convertf`, setting `outputgroup` to YES, `familyname` to NO and `hashcheck` to NO. The resulting EIGENSTRAT `ind` file had population and individual names fused in column 1, sex in column 2 and individual number in column 3 (e.g. Mapuche:GS000011196-ASM F 13199). In order to keep proper population and individual names, we sorted the `ind` file, obtaining individual ID, sex and population ID in columns 1, 2 and 3, respectively (e.g. GS000011196-ASM F Mapuche). Finally, D-statistics were run using `qpDstat` implemented in ADMIXTOOLS. The population combinations for each test are shown in **Supplementary File 1A-C**. Results were considered significant if the Z-score > |3|. Even though some populations were sequenced with more than one technology [26], we show all results in **Supplementary File 1**.

### 4.7 Admixture graph modeling

We used `qpGraph` implemented in ADMIXTOOLS [27] with default settings except the following: `useallsnps`: YES, `diag: 0.0001`, `lambdascale: 1`. We evaluated model fit based on the model’s score and the maximum |*Z*| –score, comparing predicted and observed values of the statistics. We explored two different models.

The first model explains the population splits and gene flow of Mapuche-Huilliche and the populations from Southern Cone region. We started with a suitable skeleton phylogenetic tree without admixture events and consisting of *Mbuti, Anzick1, Argentina_ArroyoSeco2_7700BP, Chile_LosRieles_5100BP, Aymara* and *Chile_Conchali_700BP*, reflecting the ideas of previous studies [6], [28], [8]. Then we successfully added a Southern Patagonia branch with *Chile_WesternArchipelago_Ayayema_5100BP, Chile_WesternArchipelago_Kaweskar_1200BP* without admixture events. Due to the affinity of Mapuche-Huilliche and Kaweskar, Mapuche and Yamana as well as observations from a previous study [8] we obtained the final result.

The second model represents the joint history of Mapuche-Huilliche and populations from the Central Andes region. We started with simple tree including *Mbuti, Anzick1, Chile_Conchali_700BP, PE_LH_Inca*. We added the remaining populations one-by-one consequently without admixture events paying attention on the score of the model and the value of the worst *f*_4_-statistics.

### 4.8 Ancestry-specific effective population size

Before estimating Mapuche, European and African chromosomal segments among admixed Chileans, we used Admixture 1.3 [63] and K=4 ancestral components to identify and exclude 592 individuals with > 1% global Aymara ancestry, obtaining a final dataset of 312 individuals. To estimate ancestry-specific N_e_, we implemented the pipeline from Browning *et al* 2018 [30]. We determined the gametic phase of each haplotype with Beagle v.5 [62]. We used RFMix [33] to estimate the local ancestry of phased genotypes. The RFMix output data was then rephased to match the original phasing. We used Refined IBD [32] with default settings together with the `merge-ibd-segments` utility to assign IBD segments longer than 2 cM to individual haplotypes. We used the `filtercolumns.jar` utility and the `adjust_npairs.py` python script to calculate the adjusted number of pairs of haplotypes. Finally, we run IBDNe (version `ibdne.07May18.6a4.jar`) [34] to estimate ancestry-specific effective population sizes.

### 4.9 Simulations of local ancestry

We performed simulations of local ancestry under a neutral scenario using the software SELAM [66], similarly as described in Vicuña et al., 2020 [37]. Briefly, we modeled a single pulse of admixture between three populations occurring *g* = 10 generations ago (1 g = 28 years [67]). This time corresponds to the time where the main pulse of admixture occurred in admixed Chileans [36]. We set mean global ancestry proportions of 0.46 Native American, 0.52 European and 0.015 African, which are the same empirical proportions estimated for our Chilean population by RFMix. We set a constant effective population size (N_e_) = 6, 000 for admixed Chileans, based on previous estimates [37]. We performed simulations using diploid (haploid) sample sizes of N = 156 (312), 312 (624), 624 (1248) and 904 (1808). We simulated 189 chromosomes, each of length 20 Morgans, in order to equate the empirical length of the human genome. We mapped the physical positions of our empirical SNPs on the simulated chromosomes. Afterwards, we calculated *T* -test *P* -values for the SNP with a maximal deviation in Native American ancestry. Variants achieving a threshold of *P* <10^−5^ were considered significant [35].

### 4.10 Variant annotations

Sequence Ontology (SO) consequence type were retrieved using the web tool VEP from Ensembl [68].

## Acknowledgements

L.V. conceived the project, designed experiments and wrote the manuscript. L.V., A.M., T.N. and A.I. analyzed the data. O.K. performed simulations. L.V and V.S. supervised the work. S.E. provided data and funding. All authors critically reviewed and accepted the final version. L.V. thanks Felipe I. Martínez from Pontificia Universidad Católica de Chile for critically revising the manuscript. The authors declare that they have no competing financial interests.

---

**Supplementary Table 1.**
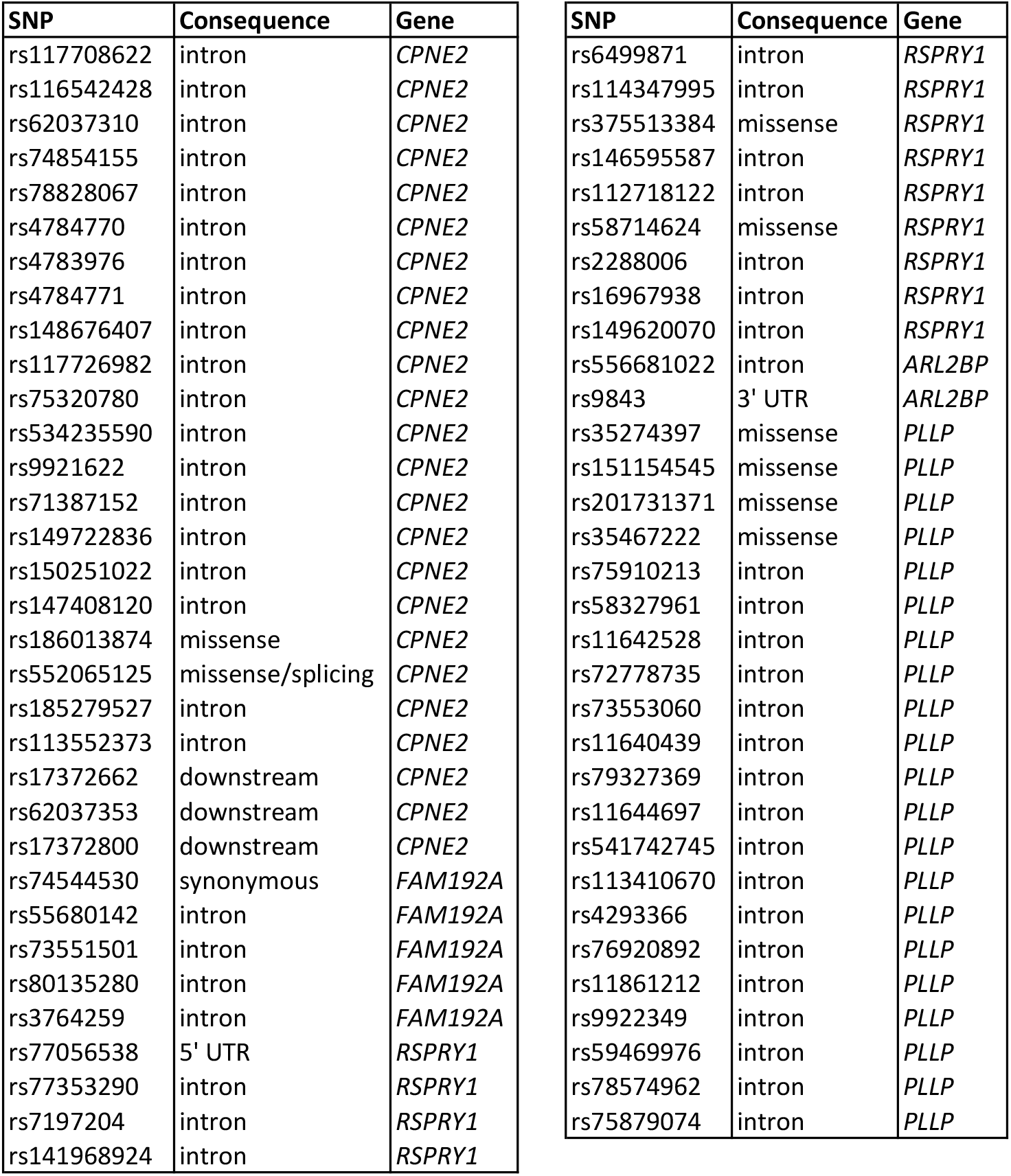
Variants undergoing Mapuche-specific post-admixture selection in admixed Chileans. Shown are the SNP ID, SO consequence type and associated gene.

